# Internal representation of hippocampal neuronal population span a time-distance continuum

**DOI:** 10.1101/475095

**Authors:** Caroline Haimerl, David Angulo-Garcia, Vincent Villette, Susanne. Reichinnek, Alessandro Torcini, Rosa Cossart, Arnaud Malvache

## Abstract

The hippocampus plays a critical role in episodic memory: the sequential representation of visited places and experienced events. This function is mirrored by hippocampal activity that self organizes into sequences of neuronal activation that integrate spatio-temporal information. What are the underlying mechanisms of such integration is still unknown. Single cell activity was recently shown to combine time and distance information; however, it remains unknown whether a degree of tuning between space and time can be defined at the network level. Here, combining daily calcium imaging of CA1 sequence dynamics in running head-fixed mice and network modeling, we show that CA1 network activity tends to represent a specific combination of space and time at any given moment, and that the degree of tuning can shift within a continuum from one day to the next. Our computational model shows that this shift in tuning can happen under the control of the external drive power. We propose that extrinsic global inputs shape the nature of spatio-temporal integration in the hippocampus at the population level depending on the task at hand, a hypothesis which may guide future experimental studies.

**Significance Statement:** The hippocampus organizes experience in sequences of events that form episodic memory. How are time and space internally computed in the hippocampus in the absence of sequential external inputs? Here we show that time and space are integrated together within the hippocampal network with different degrees of tuning across days. This was found by recording the activity of hundreds of pyramidal cells for several days. We also propose a mechanism supporting such spatio-temporal integration based on a ring attractor network model: the degree of tuning between space and time can be adjusted by modulating the power of a non-sequential external excitatory drive. In this way, the hippocampus is able to generate a spatio-temporal representation tuned to the task at hand.

## Introduction

Episodic memory holds information about spatial (where), non-spatial (what) and temporal (when) components of life experiences [1]. While spatial and non-spatial information is available in the environment (proximity to a wall, presence of a given object), temporal information is an abstract concept anchored in the dynamics of the brain. In rodents, both distance and duration have been found to be represented in the hippocampal formation, a brain area critically involved in episodic memory. This was reported in CA1 [1-5], CA3 [6-7] and the medial entorhinal cortex [8-9]. Specifically, it was shown that when rodents run in place on a wheel or a treadmill (in the absence of movement in the laboratory reference frame), hippocampal neurons fire in a sequence whose dynamics can be driven by elapsed time [2] or traveled distance [4]. While information about distance is provided by speed and self-motion cues primarily, the sequential firing of neurons in this paradigm is most likely self-organized locally at circuit-level as it occurs without any ordered arrangement of external inputs. Such internal sequences representing information relative to the past (elapsed time, traveled distance), have to be generated by an integration in time (in the mathematical sense) of spatio-temporal information. Thus, it may not be coincidence that duration and distance internal representations were observed in the same networks [4]. While mathematical models were able to reproduce the integration of time and space in different experimental paradigms [10], it remains unknown whether the same network structure can switch from encoding distance to duration.

A particularly relevant model of network structure in this background are Continuous Attractor Neural Networks (CANNs, [11-12]), which were found to generate sequences of neuronal activation from non-sequential external inputs [13-15]. In such networks, neighboring neurons within a sequence are synaptically linked, forming a circular network (or ring attractor network). They also require the presence of global feedback inhibition that allows for the firing of a few neighboring neurons at a given time (activity bump, [16]). Such type of network structure accurately describes the structure of the pyramidal layer where excitatory neurons display recurrent connections [17-18], which can in principle lead to a circular network, and interneurons provide strong feedback inhibition [19]. The main input driving sequence generation in such network models is a theta oscillation, i.e. a 6-10Hz signal present in local field potential (LFP) recordings in the hippocampus when rodents are running [20]. Wang et al 2015 [13] showed that a reduction in the power of the theta oscillation induced by silencing the medial septum led to the interruption of internal sequence generation. Therefore CANNs appear as a good candidate model to ask whether the same network organization can ground sequences encoding elapsed time and/or distance.

Here, combining data analysis of long-term sequence dynamics across days and network modeling, we asked whether the same circuit can combine and/or alternate between traveled distance and elapsed time (duration) representations and we provide a candidate mechanism by which such switching could occur. To this aim, we quantified the components of duration and distance dimensions in sequences of neuronal activation occurring in mice spontaneously running on a self-paced treadmill. Neuronal activity was sampled in CA1 using two-photon calcium imaging as previously described [5]. We found that the same network could alternate between states, sometimes combining information about duration and distance. As suggested in [6] where duration and distance related activities were equivalently found among CA1 and CA3 cells, we made the assumption that CA1 activity reflects activity in CA3 in the absence of movement in the laboratory reference frame: we thus modeled a recurrent network with feedback inhibition (CANN) driven by theta oscillation. The speed of the mouse was fed into the network through the modulation of theta oscillation power. We found that such network displays dynamics that are suitable for a dual representation of both duration and distance. Finally, by fitting experimental data, we demonstrate that the same circuit can switch between representations, under the influence of a global external signal.

## Results

We recently showed that in the absence of external constraints (no task, no reward), CA1 activity self-organizes into sequences of neuronal activation (‘run sequences’) representing distance or duration during spontaneous run episodes in head-fixed mice running on a non-motorized, uncued treadmill [5]. After expanding this dataset with additional calcium imaging experiments, we further analyzed the spatio-temporal representation in run sequences from 34 imaging sessions (n = 7 mice). In these experiments, mice were head-fixed under the objective of a microscope. Expression of calcium indicator GCaMP5, 6m or 6f was virally induced and a surgery was performed to provide a chronic optical access to the pyramidal cell layer of dorsal CA1 [5], [21]. During 20 to 30 minutes imaging sessions, 140+/-47 neurons were detected as being active within the 400×400µm^2^ field-of-view, 39+/-11% of them were activated in run sequences (Fig 1A&B, 28+/-14 run sequences per imaging session).

**Figure 1:**
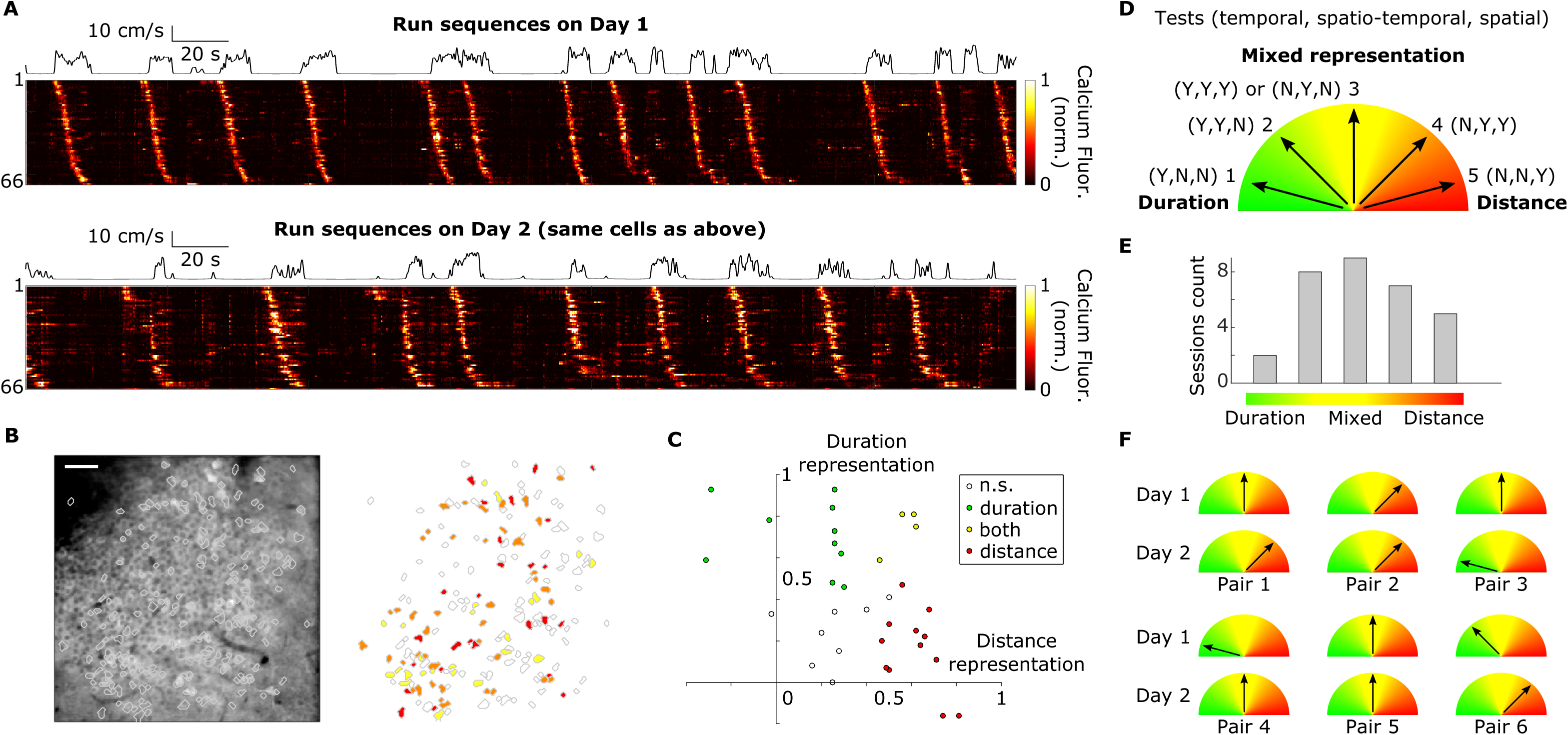
Mixed distance and duration representation in CA1. A. Calcium fluorescence) heatmap) of CA1 neurons participating to run sequences in consecutive imaging sessions; cells have been selected and ordered with respect to their activity in the first imaging session) top); black line: speed of the mouse. B. Example field-of-view in CA1 pyramidal cell layer and contours of active cells across two consecutive sessions) left) and color-coded map of cell participating in run sequences) right), red and yellow: participation on the first or second day only) respectively), orange: participation on both days. Scale bar 50µm. C. Spatio-temporal representation of the 34 imaging sessions in the distance/duration 2D space, x-axis) y-axis): Spearman correlation value between temporal) spatial) sequence slopes and running speed. D. Rule and drawing that define the five categories of spatio-temporal representation from the results of the temporal, spatio-temporal and spatial tests, Y/N) Yes or No) indicate significance) p<0.05). E. Distribution of the 34 imaging sessions across spatio-temporal representation. F. Schematics of the change in spatio-temporal representation on two consecutive days for 6 imaging session pairs.

We first applied our previously described method [5] for quantifying distance/duration representation by looking at correlations between running speed and sequence slopes in the temporal/spatial domain respectively. The network was considered as representing distance if there was a significant correlation between temporal slopes and running speeds (*spatial test*), while it was classified as representing duration if spatial slopes and running speeds were significantly correlated (*temporal test*); a network could be classified as representing both time and distance as partial correlations could be found in both tests [5]. The strength of distance/duration representation was defined as the value of the Spearman rank correlation. Imaging sessions were scattered in this 2D space (Fig. 1C), with sessions where both representations were significantly present. These results suggest that distance-only and duration-only representations are the two extreme cases of a more global, mixed spatio-temporal representation (a continuum). We next derived a new method that tests for the presence of a combination of duration and distance representations in the dynamics of run sequences (see methods and SI Appendix, Fig. S1). The basic principle is to combine the spatial and the temporal slope into a single value and test the dependence of this value with an adequate parameter that depends on speed (*spatio-temporal test*). This new test has the advantage to be less sensitive to measurement noise inherent to large-scale and fast calcium imaging. It can thus detect spatio-temporal representation where the two independent tests for distance and duration representation would not (see SI Appendix, Fig. S1D). From the results of the temporal, spatio-temporal and spatial tests, we could define five categories of representations (Fig. 1D, SI Appendix, Fig. S1) ranging from “duration-only” (case 1) to mixed forms (cases 2-4) to “distance-only” (case 5). We found that the imaging sessions that displayed a significant representation (31 out of 34 sessions) were distributed across these five categories (Fig. 1E) which confirms the capability of CA1 neurons to combine information about duration and distance [4].

We next analyzed the evolution of this representation in time, using repeated imaging sessions from the same network for two consecutive days. In 6 pairs of imaging sessions (n = 3 mice), we found that 60+/-12% of the cells participating in run sequences on the first day were recruited again in run sequences on the next day (Fig 1A&B) and that the ordered activation of these sequence-stable cells was significantly preserved (8+/-1% change in relative order, 99.9999th percentile after reshuffling). Focusing on sequence-stable cells only, we computed again the different tests for spatio-temporal representations: in 4 out of 6 paired sessions, the same set of neurons changed their dynamics with respect to speed (Fig 1F), indicating that the same network could adjust its relative representation of duration and distance.

We next designed a numerical model of the neural network to study which properties are required to enable a change in spatio-temporal integration. As discussed in the introduction,we chose a continuous attractor neural network (CANN). In order to best model a CANN grounding both duration and distance representations, we searched for specific network properties in our experimental data. As shown in Villette et al 2015, run sequences could restart where they stopped after an immobility period if the pause was shorter than 2s. In addition, run sequences could repeat one after another within continuous long run episodes. The former property suggests that there is some kind of short-term memory within the network that holds information about the last active neuron. The latter indicates the existence of functional links between first and last neurons in run sequences and supports the hypothesis of periodic boundaries in the CANN.

We designed a recurrent neural network with global feedback inhibition (CANN, Fig 2A, methods) that, when excited by a global external input, allows for the formation of a localized subgroup of firing neurons (activity bump, [11-12]). The connectivity matrix of the network integrates both local excitatory connections and a global feedback inhibition (Fig. 2B, methods). The local excitatory connections were spatially asymmetric [11], biasing the propagation of the activity bump towards one direction (Fig. 2B, methods), as observed in experimental data in which run sequences are always played in the same order. We simulated the neural activity of the network by using the following firing rate model [22]:

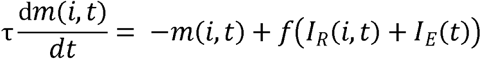

where m(i, t) is the firing rate of neuron i at time t, τ is the relaxation time constant (τ = 10ms), f is a threshold linear function, I_R_ the recurrent inputs from local excitation and global inhibition, and I£ the global external input. To account for the experimental observation that the same neurons are always recruited at the start of the run sequence, we added a bias on neurons excitability with respect to their position in the ring network (methods). A time-constant external input to all cells I£ (t) = I_0_ leads to a repetitive (circular) sequential activation of cells (Fig 2C top) that mimics experimental data. In this simplified framework, the asymmetry in the connections determines the speed of activity flow which is thus independent from the amplitude of the constant external input to the system.

**Figure 2:**
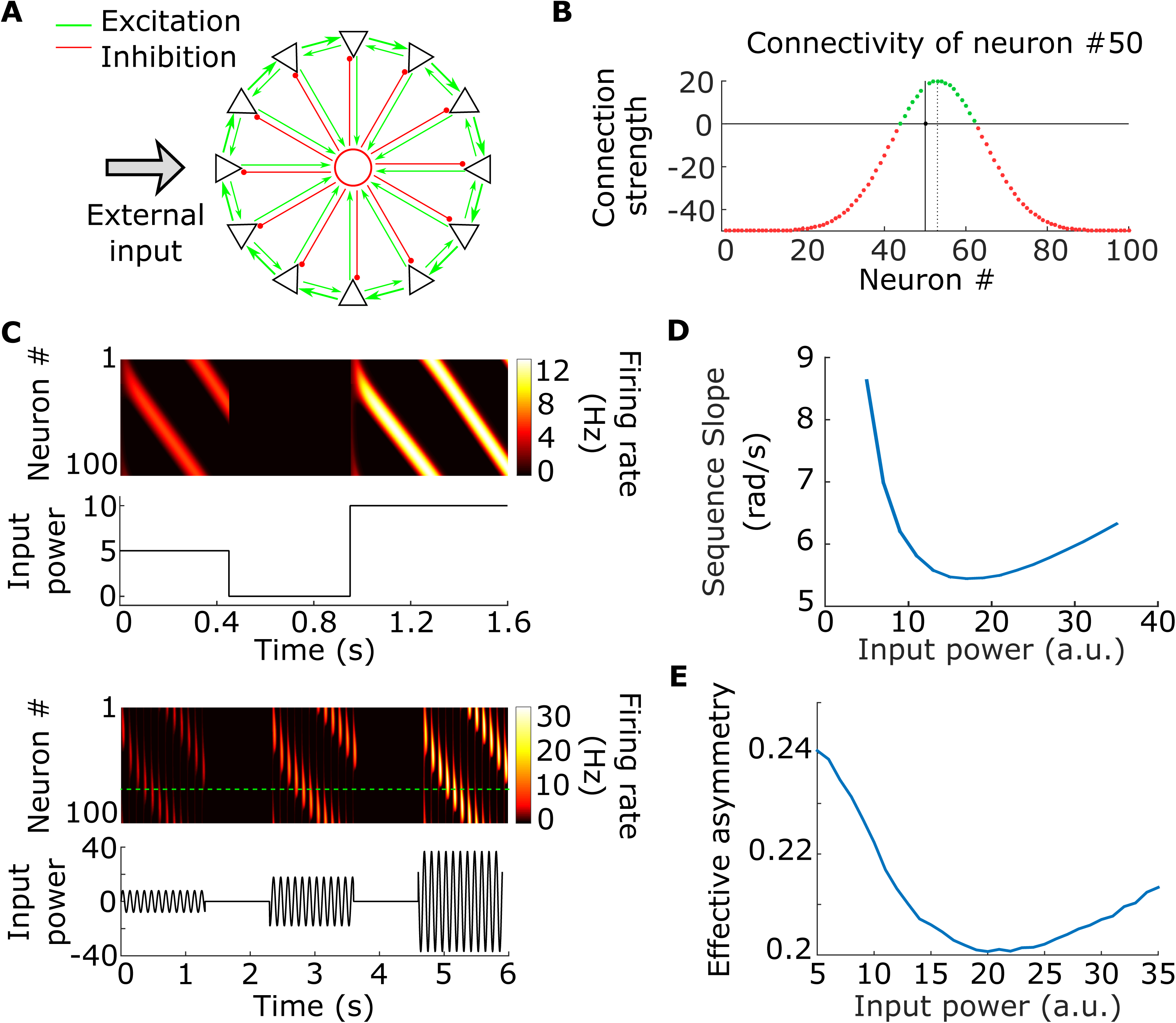
Nonlinear neural sequence dynamics in CANNs. A. Schematics of the continuous attractor network) CANNs): a ring excitatory network with global feedback inhibition. B. Total) excitatory+inhibitory) connectivity strength of neuron #50, note the asymmetry of the profile. C. Simulated neuronal sequences triggered by transient constant inputs) top) and theta input) bottom). D. Propagation velocity of the bump) or sequence slope) with respect to input power for the model with short-term plasticity and oscillatory input. E. Effective asymmetry of the network connectivity with respect to input power in the model with short-term plasticity and oscillatory input.

Following the demonstration that theta oscillations are required to generate internal sequences, the model proposed in Wang et al [13] used a similar CANN driven by an oscillatory input mimicking theta oscillation. It modulates the network activity so that at each rising phase, the most excitable cells become active, this activity bump propagates until the input power sets below threshold, at the trough, all cells are silent (Fig. 2C bottom). The group of cells active during a cycle shifts from cycle to cycle producing an overall sequence. In order to allow for the network to keep track of past activity from one cycle to the next, Wang et al.’s model [13] included synaptic facilitation and depression, which are well-known mechanisms for short-term memory in the hippocampus [23-25]. Facilitation hereby mimics the effect of residual calcium levels at presynaptic terminals that increase with firing and boost excitability, thereby creating a memory effect. Depression represents the available resources of calcium which decreases with the amount of firing and reduces excitability. We thus included theta oscillation I£ (t) = Iθ sin (2πfθ t) and synaptic facilitation and depression in our model (see methods). Their time constants were heuristically chosen to be 300ms (SI Appendix, Fig. S2). With such values, neurons reduce their firing within theta cycle bursts and active neurons remain more excitable from one cycle to the next.

In this more comprehensive model (short-term plasticity and theta input), the cycle-to-cycle velocity of bump propagation is no longer constant over different input powers but instead shows a nonlinear relationship and displays a minimum velocity (Fig. 2D). This nonlinear relationship appears both for increasing theta-modulated input or time-constant input (see Supplementary Figure 5). This is a counter-intuitive result where within a given amplitude range, increasing the amplitude of the input decreases the propagation velocity of the activity bump. Synaptic plasticities modulate the connectivity of the network and act on the propagation of the activity bump: the effective asymmetry (defined as the shift between the center of the activity bump and the center of the source term I_R_ + I£, see methods) has a similar nonlinear relationship with theta input power (Fig. 2E). Thus, the presence of short-term synaptic plasticity makes the network dynamics sensitive to external input power. This nonlinear feature is therefore a possible mechanism for the different sequence slope versus speed relationships observed in experimental data which lead to the different spatio-temporal representations (Fig 1B-D), provided that mouse speed is encoded in the amplitude of the global input and the range of amplitudes remain within a narrow window where the dependence can be assumed as linear.

To test this last hypothesis, we finally designed a speed-dependent input. Theta amplitude has been shown to increase with the running speed of an animal [26]. We thus assumed a linear relationship between theta amplitude and the speed of the mouse:

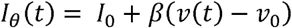

where I_0_ is the mean theta power, βthe gain applied to speed and v_0_ the median speed. The oscillatory input to the network carries speed information that will then be integrated by the CANN (Fig. 3A). Therefore, the nonlinear relationship between slope velocity and input amplitude provides a candidate mechanism for integrating time or distance in a continuous fashion through a change in input average strength. We next show that this qualitative behavior can be used to quantitatively reproduce experimental data. Firstly, we heuristically identified the span of the two-dimensional parameter space consisting of theta power I_0_ and asymmetry in connectivity δ, where values of experimental slopes could be obtained. From this phase diagram (Fig. 3B) we see that the asymmetry parameter δ sets the average value over which the slopes change, while the theta power parameter I_0_ explores the different spatio-temporal representations.

**Figure 3:**
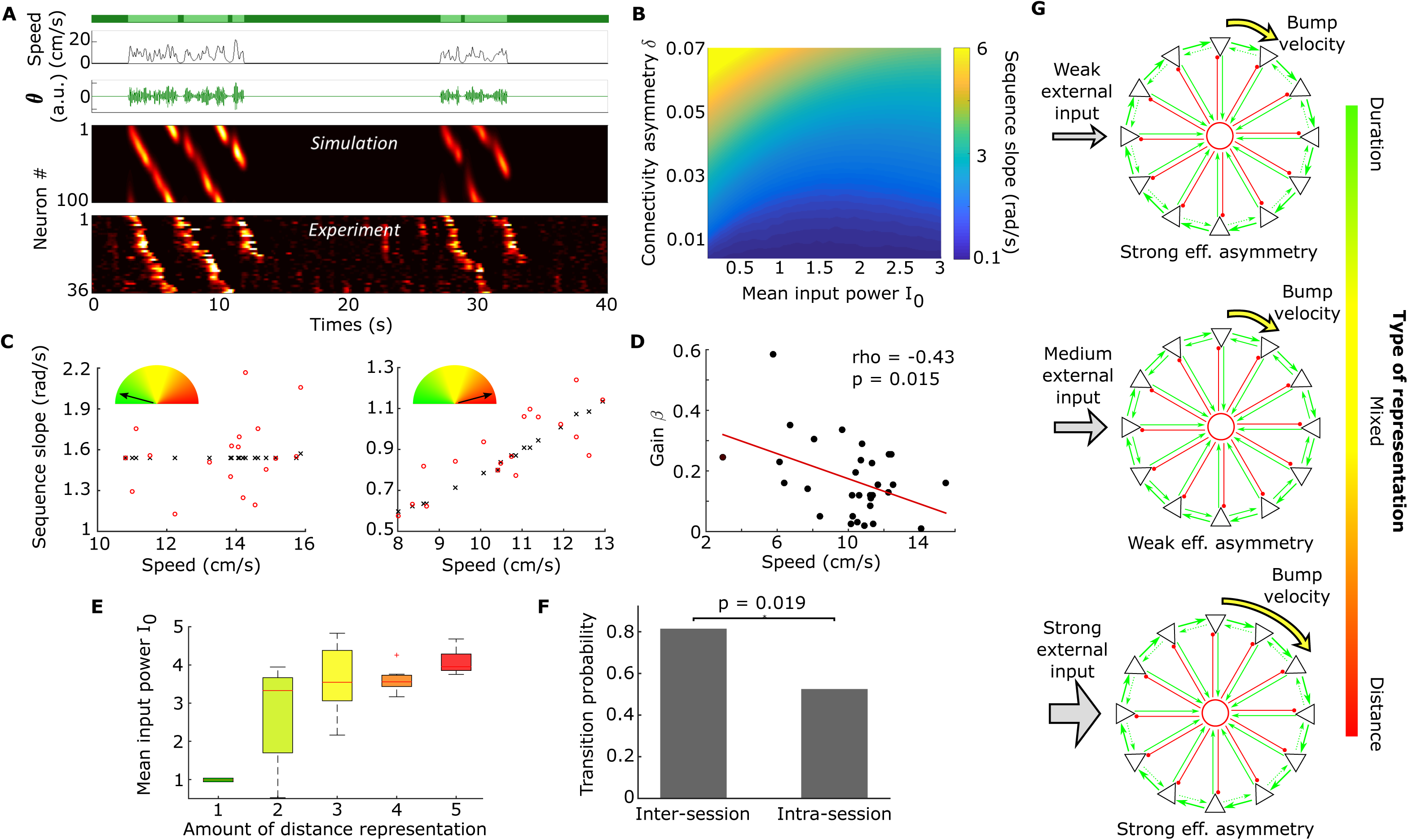
Spatio-temporal representation in CANNs. A. Realistic sequence generation: actual speed of the mouse) top, dark green: immobility period, light green: run episode), artificial theta oscillation modulated in amplitude) top middle), sequences of neuronal activation generated by the CANN) bottom middle) and actual neuronal activity recorded in CA1) bottom). B. Phase diagram in the) I_0_, δ) space with color coded slope velocities in rad/s. C. Sequence slope versus speed for duration) left) and distance) right) representation, experimental) red dots) and fitted) black crosses and lines) data. D. Scatter plot of the fitted gain parameter β and the corresponding median speed of the mouse for the 34 sessions. A linear regression shows a linear dependence between these two quantities) Pearson). E. Distribution of the mean theta power I_0_ according to the amount of distance representation showing a clear correlation) Spearman rho = 0.73, p-value = 5e-6). F. Probability for a transition in coding across sessions) inter) and within sessions) intra). A higher probability for a shift in coding is expected on a longer time scale)χ2 test, p-value < 0.05). G. Schematics of the spatio-temporal representation mechanism: for three different input powers, the network effective asymmetry and the sequence dynamics) bump velocity) change non-linearly.

A data fitting procedure was then carried out to assess the capability of the proposed model to describe real data. Indeed, the diversity of mice behavior and the range of run sequence slopes across experiments required the neural network to be robust to changes in timescale. Furthermore, as previously shown, neural sequences are not very stable from one day to the next in terms of cell participation but the cell order is mostly preserved (see above and [27]). This indicates that functional links between consecutive cells in the sequence are preserved across days. Thus, the goal was to obtain quantitative fits of the slope versus speed behavior across all data without any arbitrary adjustment of the network’s parameters. To do so, for each session we extracted the triplet of values (δ, I_0_, β that minimized the sum of squared errors between the model slopes (Fig. 3B) and the slopes of all recorded sequences during the session, as a function of the animal’s speed (see methods). To quantify the goodness of the fit, we used the Root Mean of Normalized Squared Error (RMNSE) that quantifies the mean relative deviation between data and model fit (see methods). Across 34 sessions, the fitting procedure reproduced the experimental data within an error smaller than 20%, (median RMNSE = 0.19), which is a fairly good approximation considering the purely deterministic nature of the model compared to the highly noisy data. Two example fits for a distance and duration representation are shown in Fig. 3C.

We then analyzed the relation between β and median speed v_0_ in a given session. An expected anti-correlation could be observed (Pearson rho = −0.55, p-value < 0.01, Fig. 3D). This reveals the homeostatic role of the modulatory parameter β, which upon acting on the mouse’s speed, avoids the uncontrolled increase of the input to the CANN and allows for the existence of the different spatio-temporal representations in a reduced parameter space. We finally studied whether such parameters revealed any particular feature with respect to the spatio-temporal representation as defined in Fig. 1C. I_0_ presented a correlation with the extent to which distance was represented (Spearman test, p<1e-5, Fig. 3E). This result was also expected, since “duration-only” sequences (case 1) occur when theta power is distributed around the minimum of the U-shaped function in Fig. 2D (at small inputs), while “distance-only” (case 5) are expected to occur in the right part of the U-shaped function (large inputs). Taking advantage of this correlation we quantified how the distributions of Iθ changed from one recording session to the next (inter-session) and compared to the changes from one half of a recording session to the other (intra-session). In Fig. 3F we show that the distribution of Iθ shows larger inter-session changes than intra-session changes (81% of the cases inter-session vs 52% intra-session, χ2 test, p-value < 0.05). These results suggest that the space-time representation is more likely to change on longer time scales (see SI). To conclude, we showed that a CANN with short-term synaptic plasticity has an effective asymmetry in its connectivity that nonlinearly depends on the input power (Fig 2E), it is first anti-correlated with input power for low powers, and then correlated with input power for high powers. This leads to a nonlinear dependence of the neural sequence dynamics with input power (Fig 3G). These observations demonstrate that a network integrating speed information can be externally tuned to represent any combination of space and time, and that the changes in such representation are likely to happen over long time scales.

## Discussion

The hippocampus is able to represent the diverse spatio-temporal components of experience, either mainly based on environmental inputs or on self-generated and internally computed information. Based on empirical data, we show that two internally computed information components, namely duration and distance, are embedded in the sequences of neuronal activation that occur in CA1 when a mouse is running in place on a treadmill. Instead of recruiting two distinct networks of cells that would separately integrate the temporal or spatial dimensions, we demonstrate experimentally and theoretically that the same functional circuit organization can ground both representations. This conclusion was largely enabled by our experimental paradigm that provides a wide enough temporal window on hippocampal dynamics and mouse spontaneous run behavior to cover a large enough spectrum of spatio-temporal representations.

Those representations can be generated by a continuous attractor neural network (CANN) that mimics a CA3 recurrent network. A theta oscillation modulated in amplitude by speed was used as an external input in our model so that the network could integrate speed through the propagation of an activity bump. Theta oscillations carrying speed information have been previously reported experimentally [26]; they are generated in the medial septum and directly modulate the activity of CA1 cells [28]. It has also been shown previously that when neural activity in the medial septum is pharmacologically-impaired, both theta oscillations and internal sequence generation in the hippocampus are affected [13]. This result was supported by a model of sequence generation that was the basis of our modeling work. We extended this existing framework to our experimental results by adding dependence between mouse speed and the amplitude of theta oscillation. We found that depending on the range of theta amplitude, the propagation velocity of the activity bump was either constant or linearly dependent on the speed of the mouse. Data fitting showed that these qualitative properties could be used to quantitatively reproduce experimental data. This result is explained by the presence of short-term synaptic plasticity, which dynamically influences the connectivity of the neural network. In the CANN considered here, this dynamic effect led to a nonlinear dependence of the bump propagation velocity on external input power. Even though we only considered a linear dependence between mouse speed and theta amplituden, our results can be extended to a wide set of nonlinear functions as long as a rank correlation between I_0_ and the animal’s speed v is maintained. This can be easily verified by the fact that mean theta power is the key correlate with the type of representation, therefore any function that maps the speed into I_0_ in a monotonic fashion can span the whole range of mean theta power where time-distance representation information lies.

Quantitative analysis of the data fitting showed that the type of representation (e.g. duration vs. distance) is set by adjusting the mean power of the global excitatory drive to the network. The possibility that the modulation of an external oscillatory input can change the mode of information processing in CA1 has been previously reported experimentally for gamma oscillations. Indeed, during slow gamma oscillations, CA1 activity is modulated by CA3 inputs and involved in memory processes whereas during fast gamma oscillations, CA1 is rather functionally connected to the medial entorhinal cortex, a region that transmits current spatial information [29]. Our study identifies a change in mean theta power as a possible way of shifting between spatio-temporal representations, but any global increase in excitability of the network would lead to the same effect (SI Appendix, Fig. S3). Thus, a neuromodulatory influence that changes the excitability of CA1 [30] could be another mechanism that sets the degree of distance and duration representations. Increases in theta power or in the excitability should manifest as increases in the neuronal firing rate. Future experiments, combining calcium imaging with electrophysiological measurements at the single neuron level, could test if a change in representation would be bound to a change in the firing rate, following our predictions. We also observed that the gain of the linear relationship between speed and oscillation power was inversely proportional to the median speed. This suggests a form of homeostasis that normalizes the input, a process that is expected in the hippocampal network [31] and that could be explained by the presence of feed-forward inhibition [19]. An important aspect of the fitting procedure is that it allowed us to assess the potential representation shift ability within a session, an analysis that was not possible to perform with the space-time test with experimental data due to downsampling.

Individual CA1 principal cells have been previously shown to represent both distance or duration with different degrees of tuning at single-cell level within a given recording session [4]. Here we took a different perspective by looking at the mean population dynamics (run sequence slope) of a recording session. We showed that in a given imaging session, sequential neuronal activations followed a rule for integrating space and time and that this rule could change across imaging sessions. Across all our data, the rule spanned the whole time-distance continuum, from pure time to mix to pure spatial representations. This can be explained by a model of internally generated sequences that is able to shift its coding scheme while preserving its network organization (i.e. the identity and order of cell activation in the sequence). Another relevant model of time, distance and space has been proposed by Howard et al in which a hidden layer of leaky-integrate and fire neurons with different decay time constants keep a trace of a stimuli. This first layer connects a second layer of neurons with adequate synaptic strengths in order to retrieve past events by performing a Laplace-like transform. This model nicely reproduced goal-motivated behavior. However, the model predicts scale-invariant representations (the longer the memory, the less accurate the representation). In our experimental paradigm there were no discrete stimuli: mice were running spontaneously on an empty treadmill, and we did not observe the widening of firing fields at the end of the sequence (SI Appendix, Fig. S4), the signature of scale-invariance. In the previous model, a modulation of inputs proportional to speed was used to implement path integration but this modulation was removed for simulating time representation. With our model, any type of spatio-temporal representation could be generated without the need of removing the speed-modulated input. Further work is required to combine our model of spatio-temporal representations for spontaneous behavior with the previous model for constrained behavior.

In a more general framework, CANN models for internal sequences generation can reproduce one-dimensional representations but when it comes to place cells in an open environment, it lacks a second dimension. It may be possible to extend this model to 2D using a toroidal manifold [22]. It is also possible that place cell formation goes through a different pathway that computes allocentric information [32], independent from distance and duration integration that we assume to occur in CA3. The convergence of these internal and external representations in CA1 could lead to the emergence of episodic memory: spatio-temporally ordered representation of experienced events. We indeed show that hippocampal run sequences can actually represent any type of spatio-temporal information. If an animal is trained for a given task, the hippocampus can readily adapt its integration of external inputs to represent the relevant information. For example, when a mouse is required to run for a fixed amount of time, as happens during the delay period of a spatial memory task, CA1 sequences will stretch to span and represent the entire task duration and alternatively be re-used to represent the different locations the mouse will explore in a maze [2]. This embedded sequential activity could serve as a template to temporally organize external inputs such as sensory cues or emotional events to form a memory [33]. The template has to be flexible enough in order to represent different scales and different dimensions of the ongoing experience [32]. Here we show that a recurrent network can both represent time and space, and can scale without rewiring. Future experimental work should explore this possibility.

## Materials and Methods

More details in the SI Appendix. Indicated values in the main text are mean +/- standard deviation unless otherwise stated.

All protocols were performed under the guidelines of the French National Ethics Committee for Sciences and Health report on ‘‘Ethical Principles for Animal Experimentation’’ in agreement with the European Community Directive 86/609/EEC. The experimental protocols were approved by the French National Ethics Committee under agreement #01413.03.

### Mice

Male adult wild type Swiss mice (n = 7, 30-50 g body weight) were used for experiments.

### Active cell detection

A custom-made algorithm based on PCA/ICA was used and combined to morphological identification.

### Detection of recurring activity patterns

Principal component analysis was performed on the fluorescence traces of active cells (see [5]). In order to keep a high statistical power, imaging sessions with less than 20 recurring activities (ie run sequences) were removed from analysis.

### Spatio-temporal tests

Correlations between sequence slopes in different spatio-temporal domains and mouse speed were used to test for spatio-temporal representation. In the temporal test, sequence slopes were measured in the spatial domain, in the spatial test, they were measured in the temporal domain and in the spatio-temporal test, sequence slopes were measured in a spatio-temporal domain (see more details in the SI Appendix).

### Effective asymmetry

It is defined as the shift between the bump center and the center of the source term in the neuron space.

### Firing field calculation

For each cell involved in a run sequence, we detected the onset and offset of its mean fluorescence transient over all sequences. This gave an estimate of the firing duration for each cell.

### Data sharing

Data samples can be downloaded on Figshare: https://figshare.com/s/84e04b686150c4203e67. All data and code used in analysis and for the model will be shared upon request.

## Supporting information

## Acknowledgements

This project has received funding from the ERC under the FP7 and H2020 programs (grants #242842 and 646925), the DFG (# RE 3657/1-1), the WHRI Cofund (PCOFUND-GA-2013-608765) (S.R.) and the A*MIDEX grant (No. ANR-11-IDEX-0001-02) funded by the “Programme Investissements d’Avenir” (D.A.-G. and A.T.).

