## Supplementary material for "Internal representation of hippocampal neuronal population span a time-distance continuum"

**SUPPORTING INFORMATION**

**Materials and Methods**

**Supplementary Figure 1:** Spatio-temporal tests

**Supplementary Figure 2:** Nonlinear behavior for different synaptic plasticity time constants

**Supplementary Figure 3:** Sequence dynamics changes non-linearly with excitability

**Supplementary Figure 4:** Absence of scale invariance

**Materials and Methods**

*Mice*

Male adult wild type Swiss mice (n = 7, 30-50 g body weight) were used for experiments. All mice were housed in standard conditions (12 hrs light/dark cycles, light off at 7:30 a.m., housed one per cage with water and food ad libitum). Mice were handled before recording sessions to limit head restraint-associated stress and experiments were performed during the dark cycle.

*Experimental surgery procedures*

Mice were handled before recording sessions to limit head restraint-associated stress and experiments were performed during the dark cycle. The analgesic (Buprenorphine, 0.1mg/kg) was administrated before any surgery. Viral infection was previously described [1], the virus stock solution was diluted by 1:5 (D-PBS Sigma-Aldrich) and the resultant solution was injected 2 times 200 nl of AAV2/1.Syn.GCaMP6f.WPRE.SV40 (Penn Vector Core) (AP -2.0/2.5, ML 1.6/2.1 and DV -1.3). The head-fixation bar (custom-made aluminum bar) was firmly secured with dental cement (GripCement, SuperBond, Sun Medical). Behavioral handling and imaging procedures were optimized and similarly performed as described previously [1].

*Active cell detection*

A custom-made algorithm based on PCA/ICA was used and combined to morphological identification. Movement correction was first performed for each frame using the cross-correlation with a reference image. The offset PCA method was next applied [2]. Principal components displaying a variance greater than noise were fed to an iterative ICA algorithm [3]. Cells were identified in the output of the ICA using 2D-spatial wavelet filtering matching the expected cell size (7-15µm diameter). Cell contours were next extracted. The obtained ROI were finally smoothed using a closing algorithm thus defining the active cell population. The fluorescence trace of each cell was calculated by averaging over the ROI.

*Detection of recurring activity patterns*

Principal component analysis was performed on the fluorescence traces of active cells (see [1]). GCaMP fluorescence traces were smoothed in time (Gaussian Filtering, σ=5s) before fed to the offset PCA algorithm [2]. The smoothing adds correlation between neighboring time points in order to gather the activity of cells that fire successively. The principal component which displayed recurring fluorescence patterns during each run epoch was manually selected. The derivative of that principal component was then cross-correlated with the activity of each individual cell. The correlation coefficients thus obtained were used as a metric to determine the participation of each cell to recurring activity: a threshold was calculated applying Otsu’s method [4] to the distribution of this metric in the cell population. Cells whose correlation was higher than the threshold were considered as involved in such recurring activity. In order to keep a high statistical power, imaging sessions with less than 20 recurring activities (ie run sequences) were removed from analysis.

*Sequence slope measurement*

For each recurring activity pattern, cell activation onset was defined by the maximum of the first derivative of the smoothed trace (Gaussian Filtering, σ=2s). Onsets displaying a maximal derivative smaller than 5% DF/F.s^-1^ were discarded. Cells were then ordered in a sequence according to their median onset delay in each run epoch. The sequences that involved at least half of the cells were linearly fitted using a regression algorithm that ignores outliers (Matlab function robustfit). The fits displaying a robust root mean square error exceeding 20% of the duration of the sequence were discarded. The remaining sequences were used for all quantitative analysis.

*Spatio-temporal tests*

We measured the slope of the sequential neuronal activations that occur during run with a linear fit. If done in the temporal domain (neural activity vs time), this procedure provides the temporal slopes $S_{t}$. If done in the spatial domain (neural activity vs distance), it provides the spatial slopes $S_{d}$. In the case of neural dynamics driven by time and space, these slopes follow the equations:

$$S_{t}=av+b$$

$$S_{d}=\frac{S_{t}}{v}=a+\frac{b}{v}$$

where $v$ is the speed of the mouse, $a$ in rad/cm the amount of distance representation and $b$ in rad/s the amount of delay representation. In this formalism, if $b=0$, the neural dynamics follows run distance only, if $a=0$, it follows elapsed time only, and if $a\neq0$ and $b\neq0$ the neural dynamics follows *both* spatio-temporal components.

In calcium imaging data, the slopes $S_{t}$ and $S_{d}$ are measured with an uncertainty that comes from background signal inherent to fast large-scale calcium imaging and from the detection of the activity onset in noisy cell’s fluorescence signal. Thus, relationship $S_{d}=\frac{S_{t}}{v}$ is not exact in experimental data which justifies the independent measurements of these two slopes.

In order to test for the presence of spatio-temporal representations in neural dynamics, we need to show that $a$ and/or $b$ are significantly different from zero. For this purpose, we test the Spearman correlations of $S_{t}$ versus $v$ (*spatial test*) and $S_{d}$ versus $1/v$ (*temporal test*). In the case of weak correlations in both dimensions, it is possible that none of the tests are significant but a test including both components be significant. Indeed, as noise does not sum linearly, it is relevant to combine $S_{t}$ and $S_{d}$ in order to reduce the contribution of noise. Using

$$S_{t}-v_{0} S_{d}= av-b\frac{v_{0}}{v}+b-av_{0}$$

where $v_{0}$ is the median value of the speed across all run episodes in the considered imaging session, we test the correlation of $S_{t}-v_{0} S_{d}$ versus $av-b\frac{v_{0}}{v}$ (*spatio-temporal test*). For this test, an estimate of $a$ and $b$ was required. To obtain these values, we linearly fitted the most correlated relationship among $S_{t}$ versus $v$ or $S_{d}$ versus $1/v$.

If the spatio-temporal test only is significant, or if all tests are significant, the neural dynamics follows *both* time and space. If the spatio-temporal test and only another one is significant, it is possible that the former derives directly from the latter (eg when $\alpha=0$, the temporal and the spatio-temporal tests are the identical). In such case we finally test if the increase in correlation in the spatio-temporal test is significantly higher than expected by chance by reshuffling: if the temporal (respectively spatial) and spatio-temporal tests are significant, we reshuffle $S_{t}$ ($S_{d}$) with respect to $v$ 1000 times and use the 95^th^ percentile of the correlation value on the reshuffled spatio-temporal test as a threshold for significance. With these three correlation tests and the reshuffling, we can define 5 types of spatio-temporal representations that range from pure delay (1) to mixed space and time (2-4) to pure distance (5), see Suppl. Fig. 1.

*Continuous Attractor Neural Network*

In our model, $N$ neurons are placed on a ring: the boundary conditions are periodic, all the functions defined below follow the rule:

$$G\left( i,t \right)=G\left( i+N,t \right)=G(i-N,t)$$

Where $i$ is the neuron number and $t$ is the time. The firing rate $m\left( i, t \right)$ is defined through the following set of equations:

$$\tau\frac{dm\left( i, t \right)}{dt}= -m\left( i, t \right)+f\left( I_{R}\left( i,t \right)+I_{E}\left( t \right) \right)$$

where $f$ is a linear threshold function, the threshold varies from neuron to neuron between 0 and 0.02 to simulate different degrees of excitability, leading to a stable order in the recurring sequences.

$$I_{R}\left( i,t \right)=\frac{1}{N-1}\sum_{j\neq i}^{N} W_{ij}m\left( j, t \right)U$$

$$I_{E}\left( t \right)=I$$

*I_R_* is the summed recurrent input from neighboring neurons that depends on the connection weight *W_ij_* defined as a circular Gaussian with mean shifted by an asymmetry δ (see figure 2B). Weights decrease symmetrically depending on neuron “distance” to one another in the circular arrangement.

$$W(i, j)=J_{1}\left( e^{-\left( i-j-\delta\right)^{2}}+e^{-\left( i-j-\delta-N \right)^{2}}+e^{-\left( i-j-\delta+N \right)^{2}} \right)-J_{0}$$

*U* is defined as a fixed constant (*U*=0.15) and *I* is the amplitude of a time-constant input to all neurons (*I* specified in respective graphs).

In the full model with short-term synaptic plasticity and theta oscillation as an input, new variables $x$ and $u$ to represent changes in Ca2+ available to the cell (depression, $x$) and Ca2+ accumulated inside the cell (facilitation, $u$) with respective time constants $\tau_{D}$ and $\tau_{R}$ [5].

$$\frac{dx(i, t)}{dt}=\frac{1-x\left( i, t \right)}{\tau_{D}}-u\left( i, t \right)x\left( i, t \right)m(i, t)$$

$$\frac{du(i, t)}{dt}=\frac{U-u\left( i, t \right)}{\tau_{R}}-U(1-u\left( i, t \right)m\left( i, t \right))$$

The recurrent input to the neurons is now:

$$I_{R}\left( i,t \right)=\frac{1}{N-1}\sum_{j\neq i}^{N} W_{ij}m(j, t)x(j, t)u(j, t)$$

The external input is a sinusoidal representing the occurrence of theta modulation in the hippocampus:

$$I_{E}\left( t \right)=I_{\theta}\sin(2\pi f_{\theta}t)$$

Where $I_{\theta}$ and $f_{\theta}$ are the power and frequency of the oscillation. In this study, the frequency was fixed at 9 Hz.

*Effective asymmetry*

The effective asymmetry is defined as the shift between the bump center and the center of the source term in the neuron space. It is calculated by subtracting the weighted average positions of $f\left( I_{R}\left( i,t \right)+I_{E}\left( t \right) \right)$ by the one of $m\left( i, t \right)$ at a time point $t$ where the firing rate is maximum.

*Goodness of fit*

The goodness of fit was quantified via the Root Mean of Normalized Squared Error (RMNSE), defined as follows

$$RMNSE= \sqrt{\frac{1}{N}\sum_{i} \left( \frac{y_{i}^{data}-y_{i}^{fit}}{y_{i}^{fit}} \right)^{2}}$$

This quantity measures the average local the error between the experimental data and the fitted curve. Multiplying RMNSE by 100 gives an idea of the average error in % between the measurement and the fit.

*Firing field calculation*

For each cell involved in a run sequence, we detected the onset and offset of its mean fluorescence transient over all sequences; the onset being the time when the fluorescence transient reached 10% of its maximum value and the offset the time when the trace derivative reached its minimum value. This gave an estimate of the firing duration for each cell.

*Time scale of changes in representation*

Taking advantage of the correlation between I_θ_ and the representation type, together with noiseless nature of the fitted data; we defined an estimation of the potential transition between representations over the two time scales considered in the manuscript, namely inter-session and intra-session. For the inter-session, we compared the distributions from all consecutive recording sessions. When two consecutive distributions were different (Kolmogorov-Smirnov test over 95% confidence level), a potential transition of representation was considered. The number of potential transitions divided by the total number of possible transitions defined the inter-session transition probability. For the intra-session case we split the data for each recording session in two halves and compared the differences in the distributions of Iθ in these two halves. Whenever the two distributions of Iθ were significantly different, we assumed a potential transition within the session. The total number of successful tests divided by the total number of recording sessions defined the intra-session transition probability. We found significantly smaller probabilities of intra-session potential transitions with respect to the inter-session counterpart (52% vs 81%, χ2 test, p-value < 0.05). We have used the term “potential transition” to indicate that the representation shift within a session, based only on the I_θ_ distributions, does not guarantee an actual shift in the representation (see for example the overlap between type 3 and type 4 representation in Fig. 3E). This means that the effective percentage of transitions within a session is likely to be even smaller than those reported here, and our estimation should be regarded as an upper bound.

*Dependence of representation with behavioral input*

We tested the possibility that shifts in representation could be related to changes in the distributions of the only behavioral input into the model (animal’s speed). To do this we compared the inter-session speed distributions (with sessions usually separated by one day or more) via a Kolmogorov-Smirnov test with 95% confidence level. The percentage of times that a representation change was accompanied by a difference in speed distribution was not significantly different from the percentage where a change in representation occurred in the absence of it (66% vs 60%). This allowed us to conclude that changes in the external/behavioral input variables are unlikely to be the at the basis of representation shifts.

*Stability of representation before/after a running break*

We further tested the stability of distance vs duration representation before/after a break in running. For this we applied our spatio-temporal tests on the first and second halves of sequences successively. By definition, the first halves of sequences followed while the second halves preceded an immobility period. Because fitting only half of the sequences led to a worsened accuracy of slope estimation, only 18 out 34 sessions displayed a significant spatio-temporal representation on both sequence halves. Among them, 56% had the same representation on both halves, 17% switched to distance representation and 28% to time representation. With such results, we can now conclude that the stability of spatio-temporal representation type is not influenced by a preceding immobility period.

**Supplementary Figures**

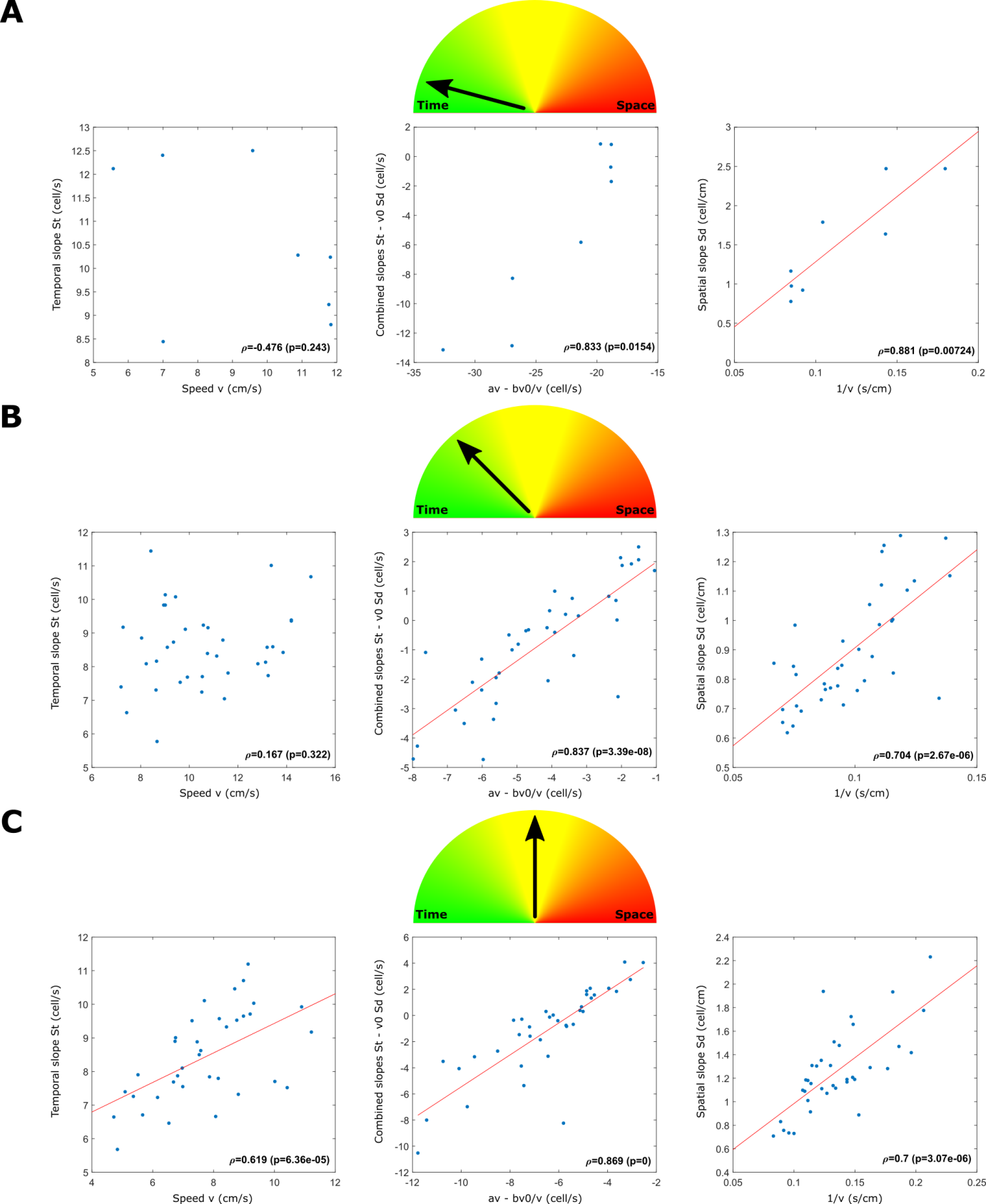

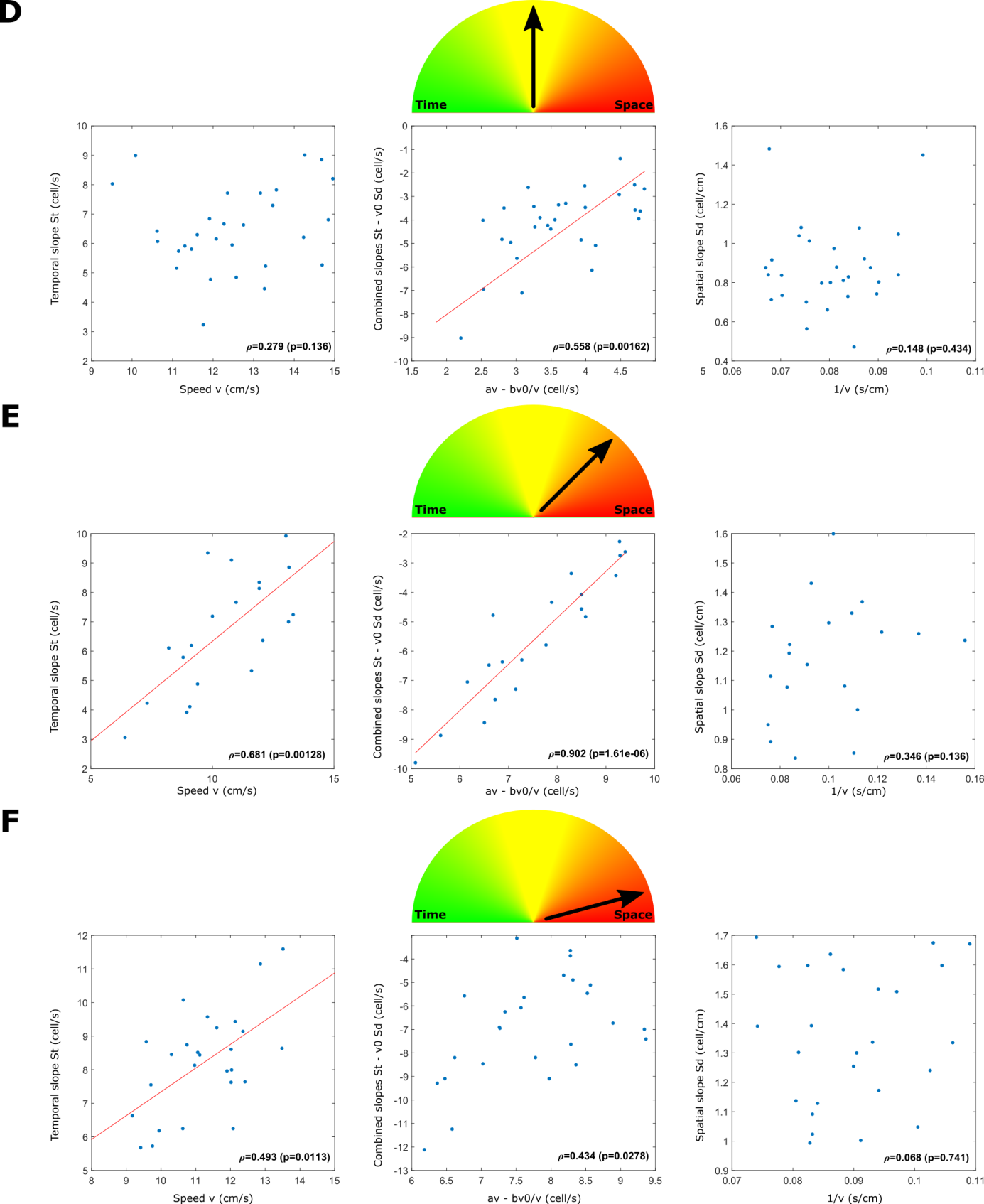

**Supplementary Figure 1:** Spatio-temporal tests

The 6 cases of spatio-temporal representations observed in data. For each test, the strength ρ and the p-value of the Spearman correlation is indicated. A. Case 1 “Pure time representation”, B Case 2, C Case 3 with the three tests that are significant, D Case 3 with the spatio-temporal test only that is significant, E Case 4, F Case 5 “pure distance representation”

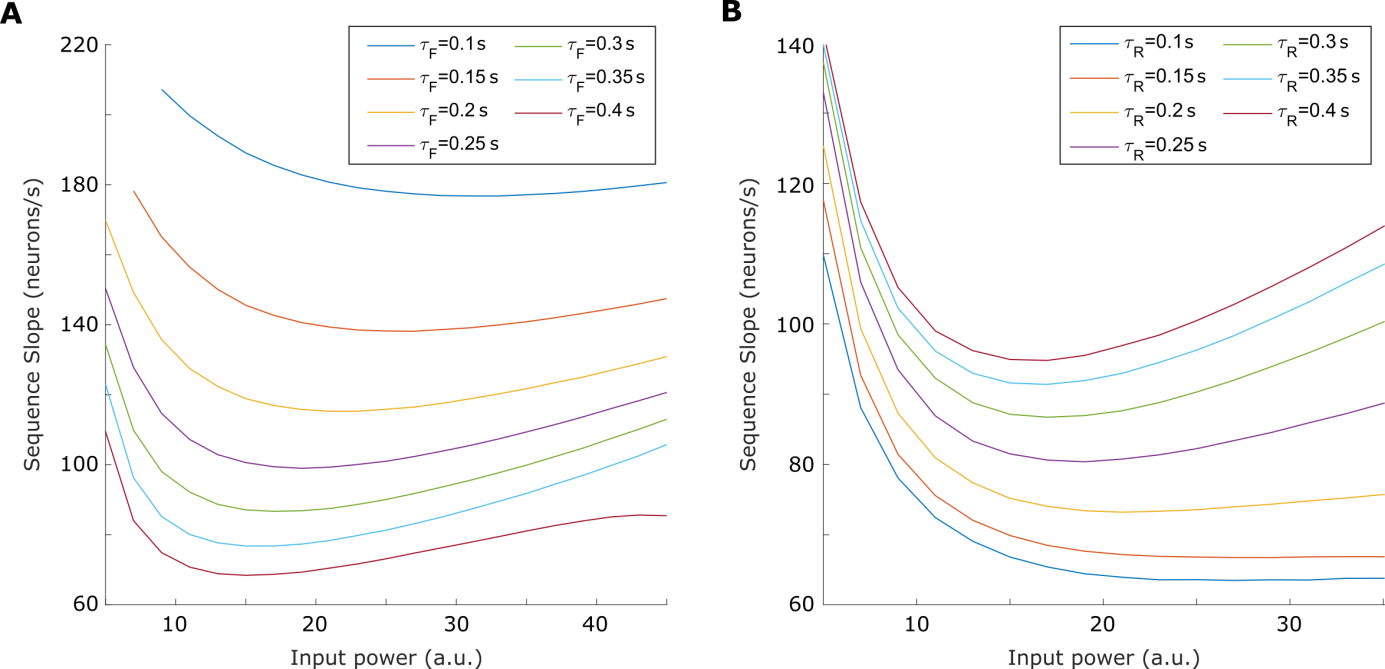

**Supplementary Figure 2:** Nonlinear behavior for different synaptic plasticity time constants. A. Facilitation time constant from 0.1 to 0.4 s, B Depression time constant from 0.1 to 0.4s.

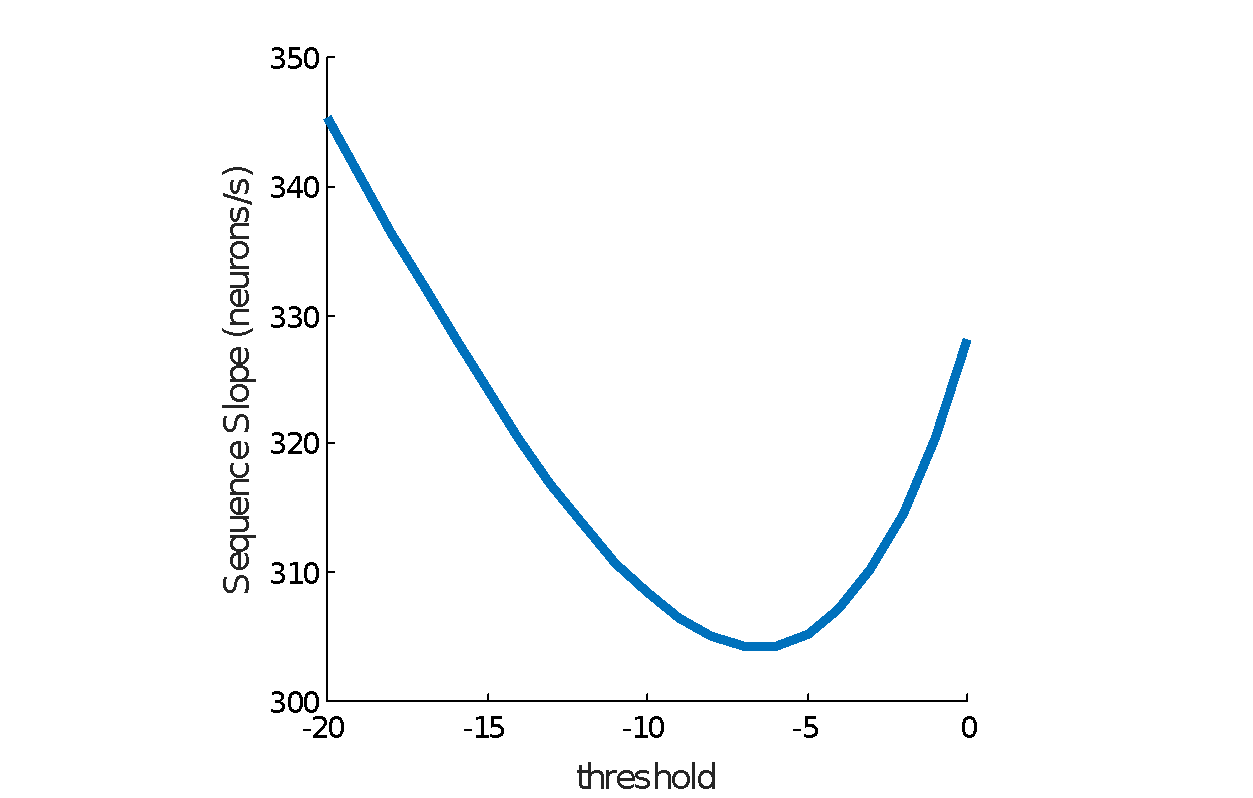

**Supplementary Figure 3:** Sequence dynamics changes non-linearly with excitability. Sequence slope was measured in successive simulations with different spike threshold in the linear threshold function *f*. It is an equivalent of a general excitability change in the network. The curve has the same features as the one in Fig 2D: it is nonlinear with a minimum velocity.

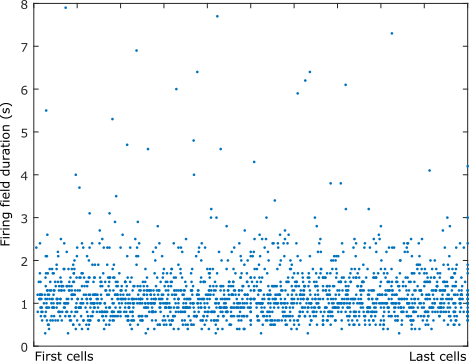

**Supplementary Figure 4:** Absence of scale invariance

Firing field calculated for the 1888 sequence cells across 34 imaging sessions with respect to their position in the sequence. Firing fields were defined as the duration between the onset and the maximum of the calcium transient (see Villette et al. 2015). No significant increase was observed (p > 0.05, spearman correlation test).

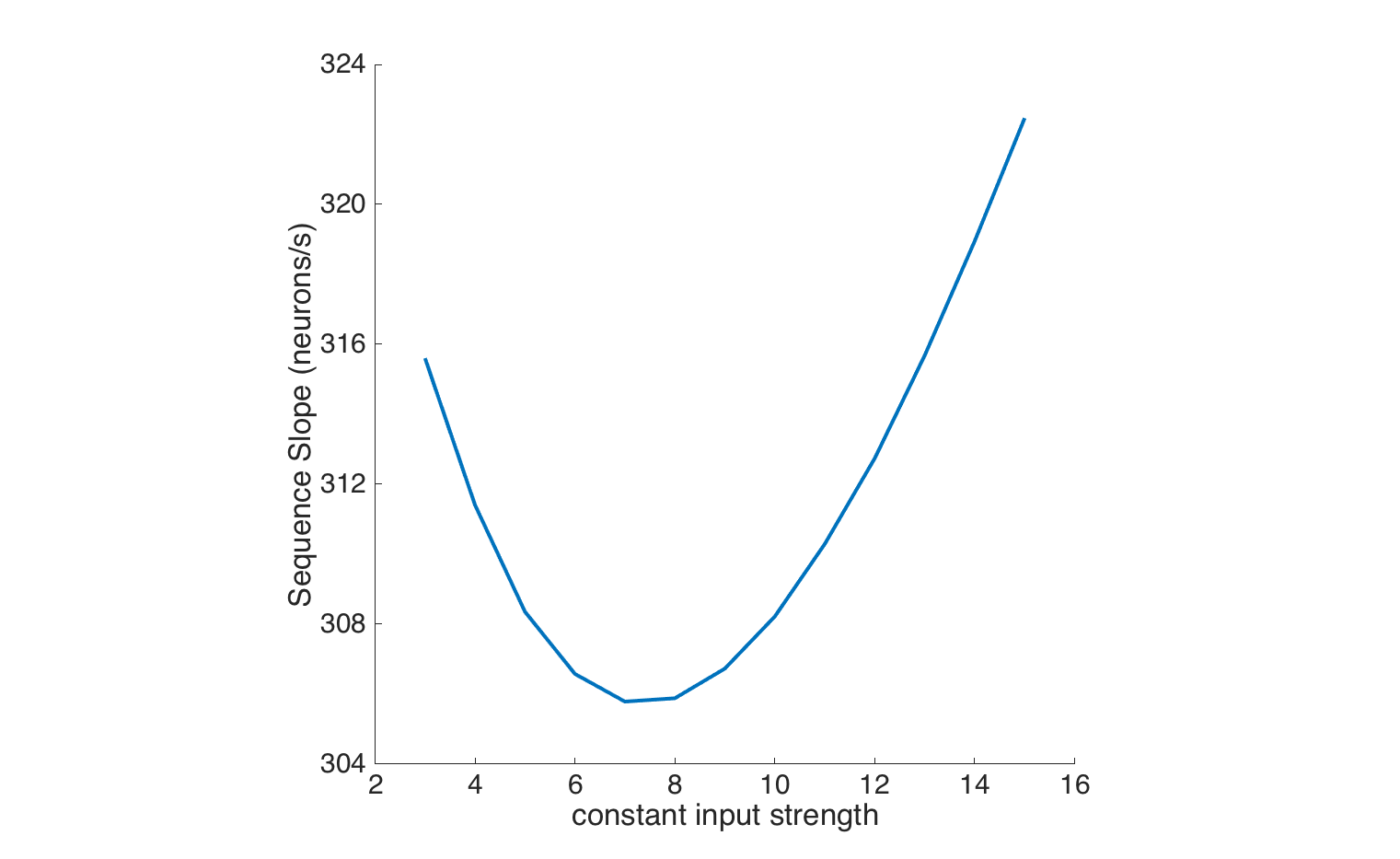

**Supplementary Figure 5:** Sequence dynamics change non-linearly with time-constant input strength. Here we simulate a version of the model that includes short-term plasticity but where the input is constant in time instead of theta-modulated. The curve is qualitatively similar to the one achieved with theta-modulated input.
