## Supplementary figures and images for "Internal representation of hippocampal neuronal population span a time-distance continuum"

### Supplementary file 2

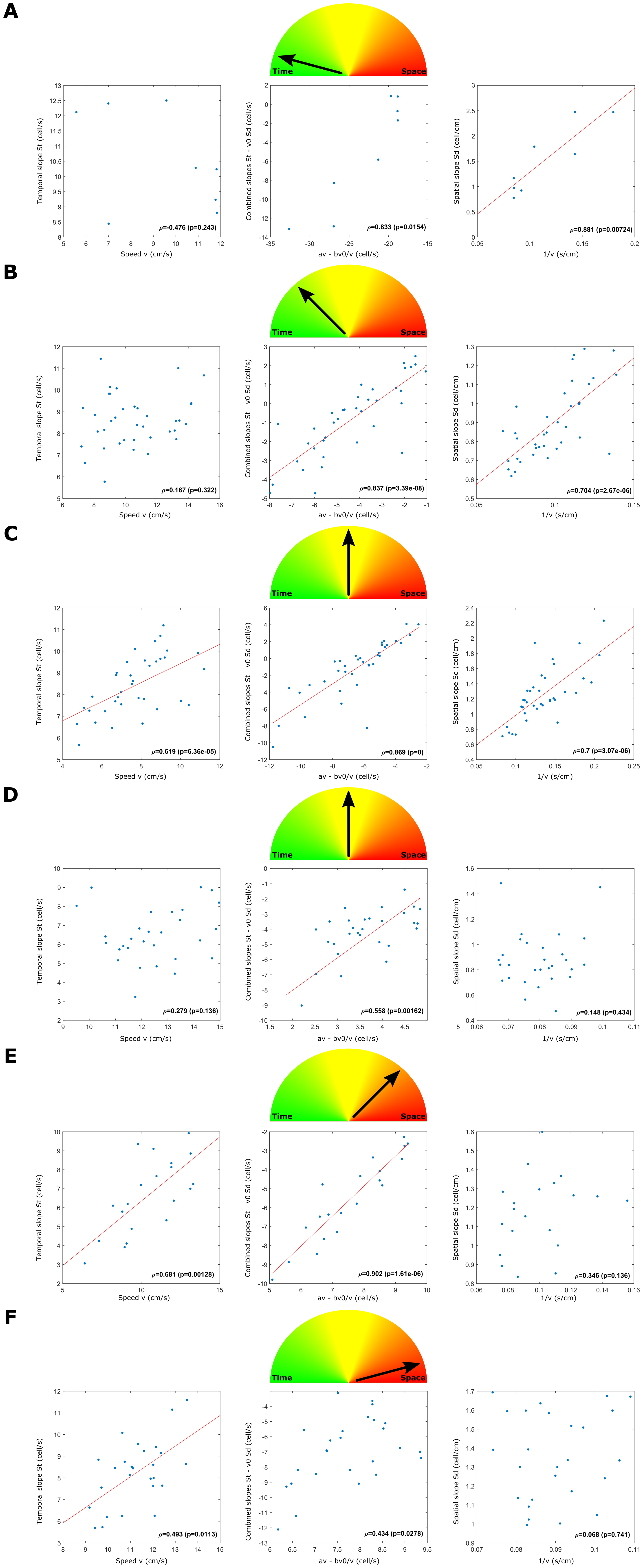

### Supplementary file 3

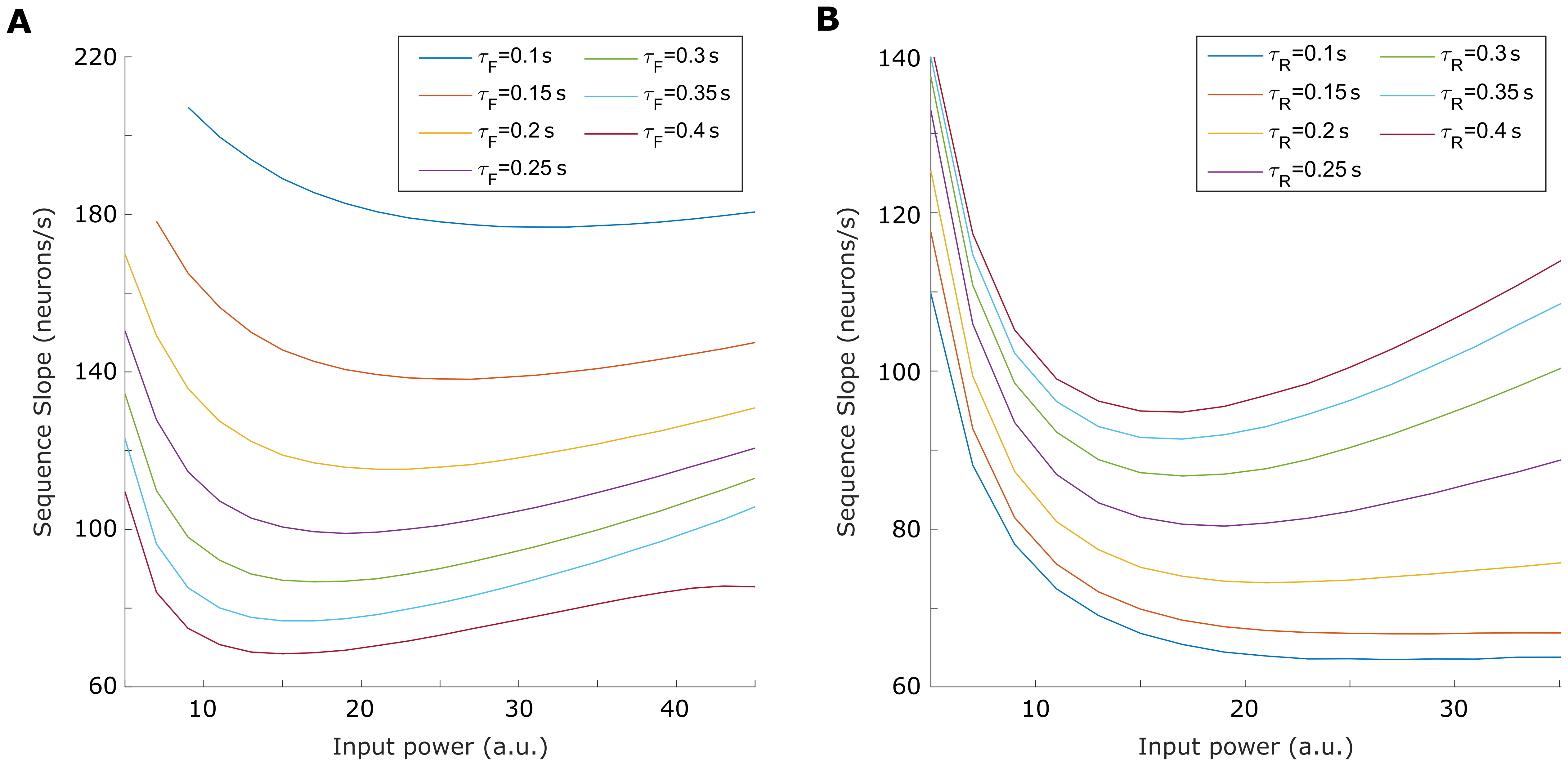

### Supplementary file 4

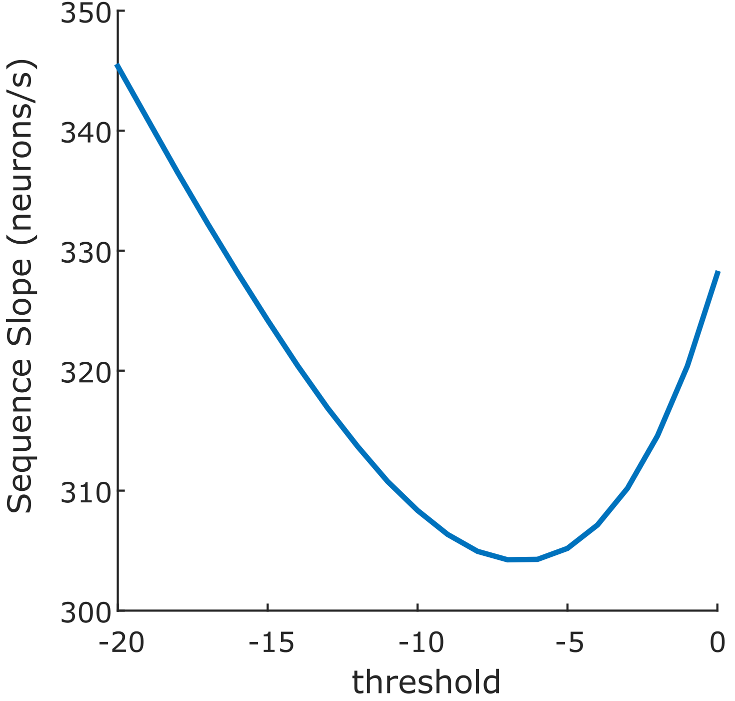

### Supplementary file 5

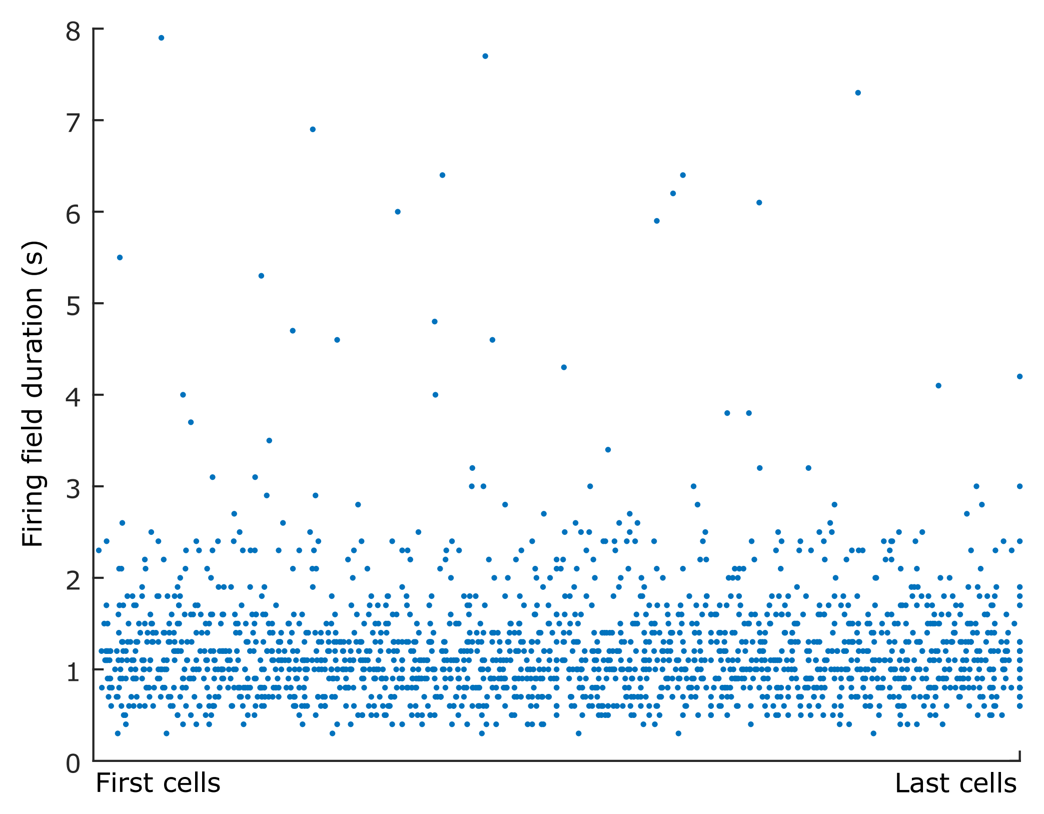
